# Extensive variation in the mouse intelectin gene family: recent duplications, deletions and inactivating variants result in diversity in laboratory strains

**DOI:** 10.1101/2021.03.31.437930

**Authors:** Faisal Almalki, Eric B. Nonnecke, Patricia A. Castillo, Alex Bevin-Holder, Kristian K. Ullrich, Bo Lönnerdal, Linda Odenthal-Hesse, Charles L. Bevins, Edward J. Hollox

## Abstract

Intelectins are a family of multimeric secreted proteins that bind microbe-specific glycans. Both genetic and functional studies have suggested that intelectins have an important role in innate immunity and are involved in the etiology of various human diseases, including inflammatory bowel disease. Experiments investigating the role of intelectins in human disease using mouse models are limited by the fact that there is not a clear one-to-one relationship between intelectin genes in humans and mice, and that the number of intelectin genes varies between different mouse strains. In this study we show by gene sequence and gene expression analysis that human intelectin-1 (*ITLN1*) has multiple orthologues in mice, including a functional homologue Itln1; however, human intelectin-2 has no such orthologue or homologue. We confirm that all sub-strains of the C57-line have a large deletion resulting in retention of only one intelectin gene, *Itln1*. The majority of laboratory strains have a full complement of six intelectin genes, except wild-derived CAST, SPRET, SKIVE, MOLF and PANCEVO, which are derived from different mouse species/subspecies and encode different complements of intelectin genes. In wild mice, intelectin deletions are polymorphic in *Mus musculus castaneus* and *Mus musculus domesticus*. Further sequence analysis shows that *Itln3* and *Itln5* are polymorphic pseudogenes due to premature truncating mutations, and that mouse *Itln1* has undergone recent adaptive evolution. Taken together, our study shows extensive diversity in intelectin genes in both laboratory and wild-mice, suggesting a pattern of birth-and-death evolution. In addition, our data provide a foundation for further experimental investigation of the role of intelectins in disease.

## Introduction

Intelectins (<u>*inte*</u>stinal <u>*lectins*</u>) are a family of calcium-dependent multimeric, secreted proteins that selectively bind microbial carbohydrates (1–3). In mammals, intelectins were initially identified in the mouse small intestine, but are found throughout vertebrates, and have a variety of roles including host-microbe interaction (3–9). In humans, there are two intelectin proteins intelectin-1 and intelectin-2, encoded by the genes *ITLN1* and *ITLN2* respectively, which are tandemly arranged on chromosome 1q23.3. Human ITLN1, also known as omentin-1, is expressed in visceral adipose as well as the intestine (1,10–12). ITLN1 binds to exocyclic vicinal 1,2-diols, a chemical moiety present in microbial carbohydrate-containing structures such as β-D-galactofuranose, which is a galactose isomer synthesized by microorganisms, including protozoa, fungi, and bacteria, but not by mammalian cells (5). The tissue expression and lectin binding properties of ITLN2 appear different (13), but are yet incompletely delineated. Intelectins have been implicated in several diseases including inflammatory bowel disease (IBD), obesity, non-insulin-dependent diabetes mellitus, and asthma (4,12,13,8).

Expression in the small intestine, and the ability to recognize bacteria-specific glycans suggests that ITLN1 has a key role in the innate immune response in the gut. Furthermore, genome-wide association studies (GWAS) have identified a number of associations with inflammatory bowel diseases. In particular, an early GWAS identified a common single nucleotide variant allele rs2274910-C associated with increased risk of Crohn’s disease (14,15). Subsequent GWAS have identified other variants within and surrounding *ITLN1* associated with Crohn’s disease and ulcerative colitis (16–18). It is likely that these associations are driven by the same causative variant, but that causative variant has not yet been identified.

The link between intelectins and disease such as IBD suggests it will be informative to pursue further research on intelectin function *in vivo* using animal models. Nevertheless, analysis of mouse intelectins is challenging as mice show strain-specific variation in intelectin gene copy number and tissue-specific expression patterns. The laboratory mouse strain 129S7/Sy encodes six intelectin genes on chromosome 1, generated by recurrent inversion and duplication, whereas the C57BL/6J sub-strain encodes a single intelectin gene, *Itln1*, remaining from a large 420 kb deletion (5). The assignment of particular mouse intelectin genes as orthologues of human *ITLN1* remains unclear.

We therefore considered it important to fully characterise the intelectin gene locus in both laboratory strains and wild mice, and investigate the complement of intelectin genes across these mice, as well as sequence and expression variation. We characterised expression patterns of the intelectin genes in different tissues of different strains, with a particular focus on the gastrointestinal tract, where intelectins appear to be commonly expressed in vertebrates (3). This characterisation will provide a firm grounding for experiments using mouse models to identify or study intelectins in gastrointestinal, lung, metabolic, and infectious diseases. Importantly, because analyses of the genomes of laboratory mice and wild mice relied on mapping to a reference genome from the C57BL/6J sub-strain, all members of the the intelectin family except *Itln1* appear to be absent from these genomes. In 15 laboratory strains and 29 wild mice we mapped publicly-available short read data to a sequence contig of the intelectin gene region derived previously from a 129S7/Sv strain mouse. We confirmed and characterised the deletion in C57BL/6J, show it is present also in the three progenitor strains (i.e., C58/J, C57L/J, C57BR/cdJ) for C57-lines, and found novel deletions in both wild and laboratory strains of mice. Moreover, we characterized the sequence variation of intelectin genes, in the context of the repeated nature of this region.

## Results

### Human ITLN1, but not ITLN2, has multiple orthologues in mice

We selected 20 coding sequences of representative intelectin genes from six primates and nine rodents. A phylogenetic tree shows that Muridae rodents have one intelectin gene, with the exception of mice where a recent burst of duplications has generated six intelectin genes as previously noted (8) (Figure 1a). *Itln1*, *Itln2*, *Itln5* and *Itln6* are annotated as full-length proteins of 313 amino acids. Protein products from *Itln3* and *Itln4* genes are less clear, and indeed *Itln3* is annotated with an early truncated mutation predicting a foreshortened peptide of 142 amino acids. Nevertheless, both *Itln3* and *Itln4* genes exhibit very high sequence identity to the other mouse intelectin genes, therefore have arisen by very recent duplication.

**Figure 1.**
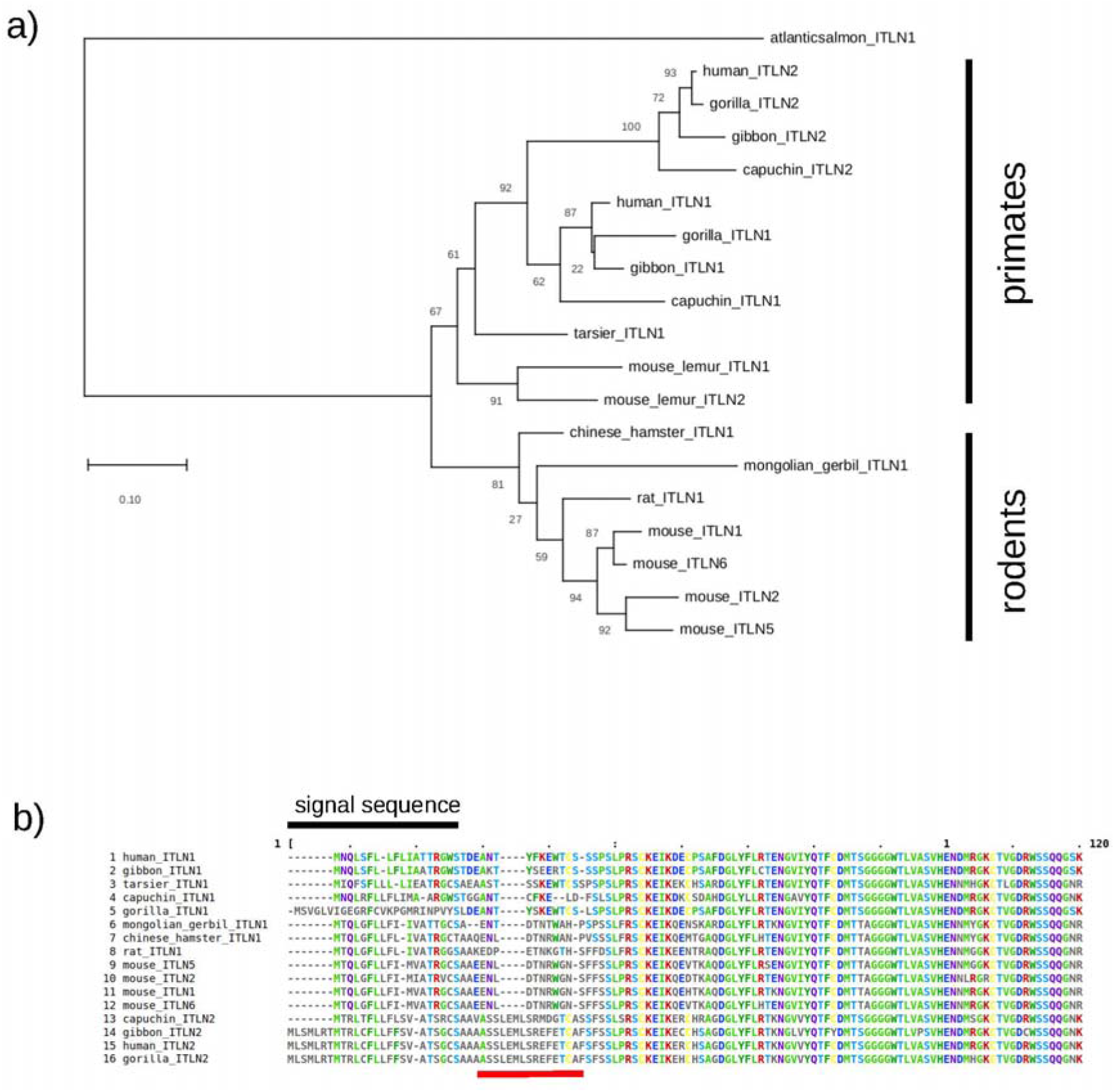
Intelectin genes in rodents and primates. a) A phylogenetic tree based on amino acid sequences is drawn to scale, with branch lengths measured in the number of substitutions per site. Numbers at each node indicate bootstrap support for that node. b) Amino acid sequence alignment of the N-terminal region of intelectin.

In primates, an independent duplication event has generated two intelectin genes, *ITLN1* and *ITLN2*, encoding proteins of 313 and 325 amino acids, respectively. This event may have occurred early in primate evolution, with the tree suggesting a duplication before the divergence of lemurs and other primates, followed by loss of *ITLN2* in lemurs and tarsiers. In the lemur lineage, an independent duplication of *ITLN1* has generated another intelectin gene, named *ITLN2,* though very distinct from other primate *ITLN2* sequences. From the phylogenetic analysis it is clear that there is not a simple one-to-one relationship between human and mouse intelectin genes. In addition to length, inspection of the N-terminal protein sequence alignment (Figure 1b) confirms that ITLN2-like proteins are distinguished from ITLN1-like proteins by this distinct sequence of amino acids at the N-terminus of the mature peptide, and shows that all mouse intelectins are ITLN1-like, with no orthologue of human ITLN2.

We addressed the possibility that a functional homologue – an intelectin that is not necessarily orthologous but fulfils the same function in human and mouse – exists for both human genes. To explore this, we noted the constitutive expression pattern of ITLN1 and ITLN2 in humans and examined the pattern of expression of the intelectin genes in the intestine of 129S2/SvPasCrl mice, which, like 129S7/Sy, carry the full complement of six intelectin genes (Supplementary table 2). Inspection of the Gtex, FANTOM5 and Human Protein atlas expression data show that human ITLN1 and ITLN2 have distinct expression patterns (Supplementary figure 1). Across the datasets, for the intestine, ITLN1 is expressed in both small intestine and colon, while ITLN2 expression appears narrower.

In the mouse, we confirmed that *Itln1* in the C57BL/6Crl sub-strain is primarily expressed in the ileum, and, to a lesser extent, in the colon (Figure 2a) using RT-qPCR. In 129S2/SvPasCrl, which has the full complement of intelectin genes, simple RT-qPCR is useful to detect tissue expression patterns (Figure 2b) but is unable to unambiguously distinguish the transcripts from the individual intelectin genes due to high sequence similarity across the intelectin genes. Therefore we used a complementary strategy using high-throughput sequencing (Supplementary figure 2) to determine which orthologues were expressed in the small intestine and colon of the 129S2/SvPasCrl mice. In the small intestine we observed *Itln1* expression consistent with previous reports (4,6), but also detected trace levels of *Itln2* and *Itln6* transcripts (0.22% and 0.15% of total intelectin transcripts, respectively) (Supplementary Table 1). In the colon we observed *Itln6* expression (99.94%) and trace levels of *Itln2* (0.06%), however neither *Itln1* nor any other orthologue was detected (Supplementary Table 1). Despite the aforementioned genetic differences in encoded intelectin genes between C57 and 129 strains, the highest mRNA expression levels in mice appear to be in the small intestine, whereas extra-intestinal baseline expression is at much lower levels (Figure 2b) – a contrast to human *ITLN1* that is highly expressed in other tissues, including visceral adipose. Higher expression levels in mouse esophagus, stomach, ovary and uterus (Figure 2b) may be due to other intelectin genes being expressed in those tissues.

**Figure 2.**
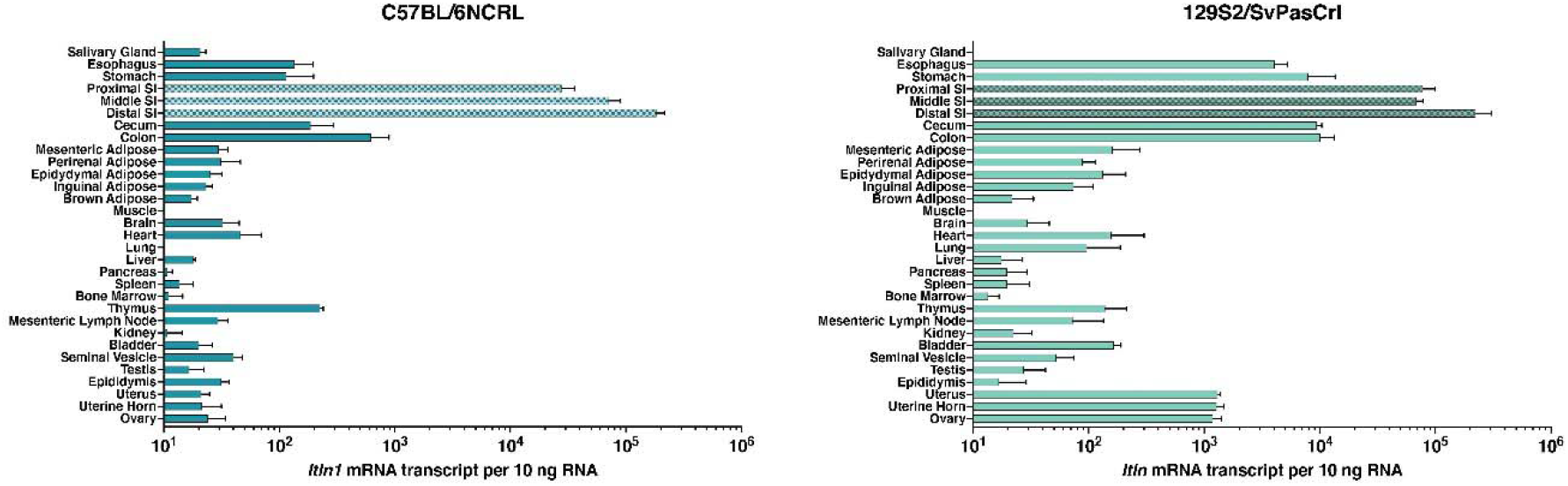
Analysis of intelectin expression in mice. mRNA expression levels measured by RT-qPCR are shown across tissues for (a) *Itln1* in C57BL/6NCRL and (b) all intelectins in 129S2/SvPasCRL. Absolute quantification of mouse *Itln* mRNA transcript counts from tissue samples (n=4 mice) determined from standard curves using a sequence specific plasmid and presented as transcripts per 10 ng total RNA. Note that qPCR primers were designed to amplify all six intelectin transcripts from 129S2/SvPasCRL with equal efficiency. Error bars represent standard error of the mean.

Looking across other mouse strains, we confirmed a higher level of intelectin expression in ileum compared to colon (Figure 3). Using the next generation sequencing strategy described above, we analyzed the intelectin transcripts in wild-derived strains PANCEVO/EiJ and SKIVE/EiJ and confirmed that as in 129S2/SvPasCrl only *Itln6*, and not *Itln1*, was expressed in the colon, with the small intestine expressing mostly *Itln1*, although in these cases about 20% of the intelectin transcript was *Itln6*. This strongly suggests that, after duplication, *Itln6* and *Itln1* adapted to become more tissue specific, and that the subsequent derived deletion in C57-lines removed this specificity, and resulted in *Itln1* being expressed in both tissues again – perhaps by removing key cis-acting elements.

**Figure 3.**
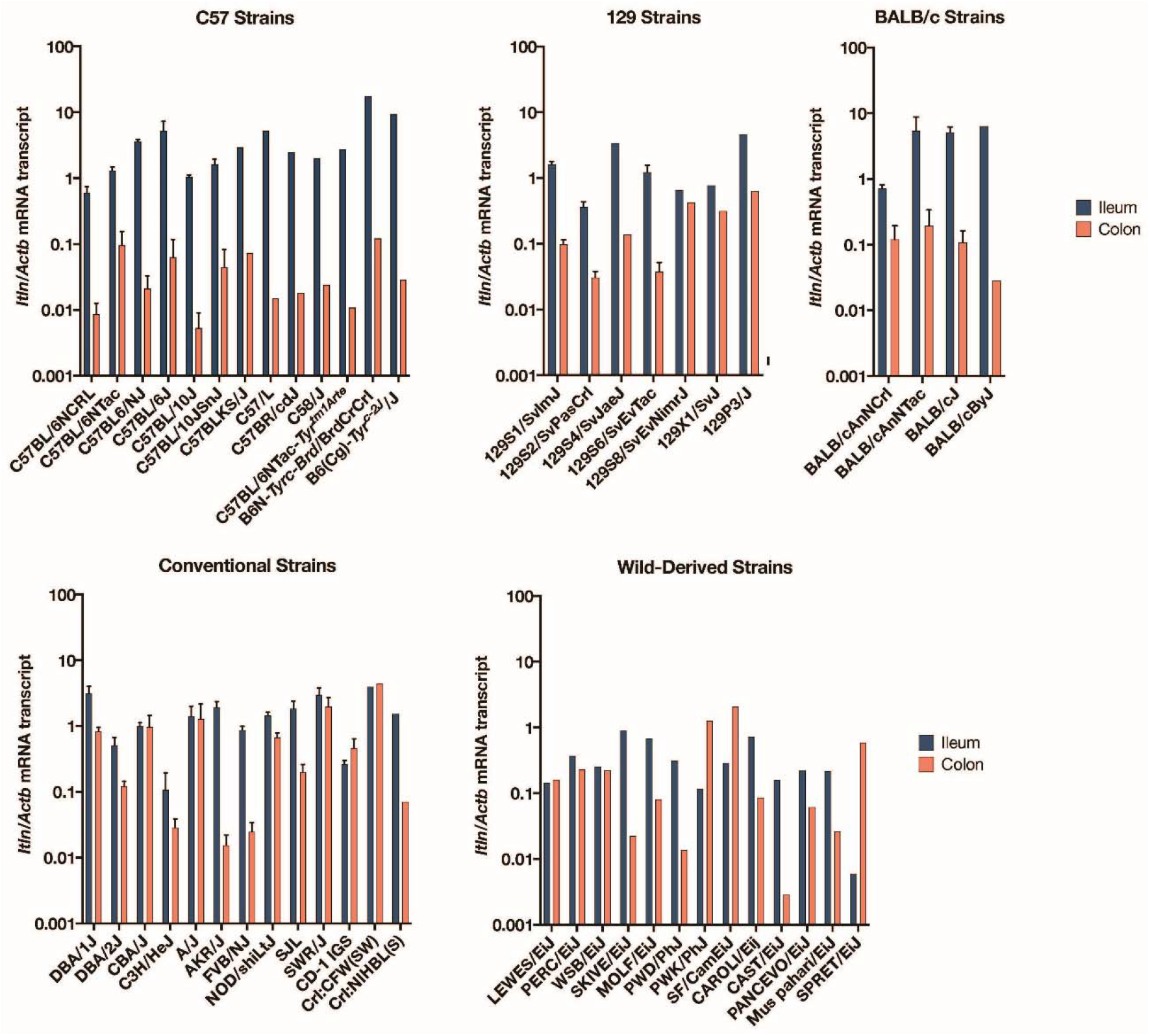
Analysis of intelectin expression in ileum and colon across different mouse strains. Quantification of mouse *Itln* mRNA levels in small intestinal ileum and colon tissue of mouse strains determined by RT-qPCR. Note that qPCR primers correspond to targets of identical sequence in *Itln1* in C57BL/6NCRL and all six intelectin mRNA in 129S2/SvPasCRL. Data are normalised to *Actb* transcript levels. Error bars represent standard error of the mean (n=4 mice). Because of high cost of some strains, specimens from a single mouse was analyzed and data presented without error bars.

### Mouse intelectin genes show extensive copy number variation

To detect whether all sub-strains of C57-lines also only had *Itln1*, we designed several paralogue ratio tests (PRTs) to test for the presence of *Itln1*, *Itln2*, *Itln4* and *Itln6* genes in mouse genomic DNA (Figure 4). Analysis of twelve C57BL/6 substrains showed that these, together with the C57 progenitor strains (i.e., C57L/J, C57BR/cdJ, and C58/J), only have *Itln1* but not *Itln2*, *Itln4* or *Itln6*, suggesting they share the same deletion (Supplementary table 2). Deletion-specific PCR primers were designed across the deletion breakpoint in C57BL/6J, which yielded PCR amplification products of identical size in the strains with the deletion. Sanger sequencing confirmed that the breakpoint was identical in these strains (chr1:173447174-173447330), within a SINE, and therefore identical by descent. Other laboratory mouse strains derived from *Mus musculus musculus* and *Mus musculus domesticus* showed the full complement of six intelectin genes, according to our PRT analyses (Supplementary table 2). We confirmed the presence of the full region in PWD/PhJ, an inbred mouse strain of the subspecies *M. m. musculus*, and deletion in C57BL/6J, using optical mapping (Figure 5).

**Figure 4.**
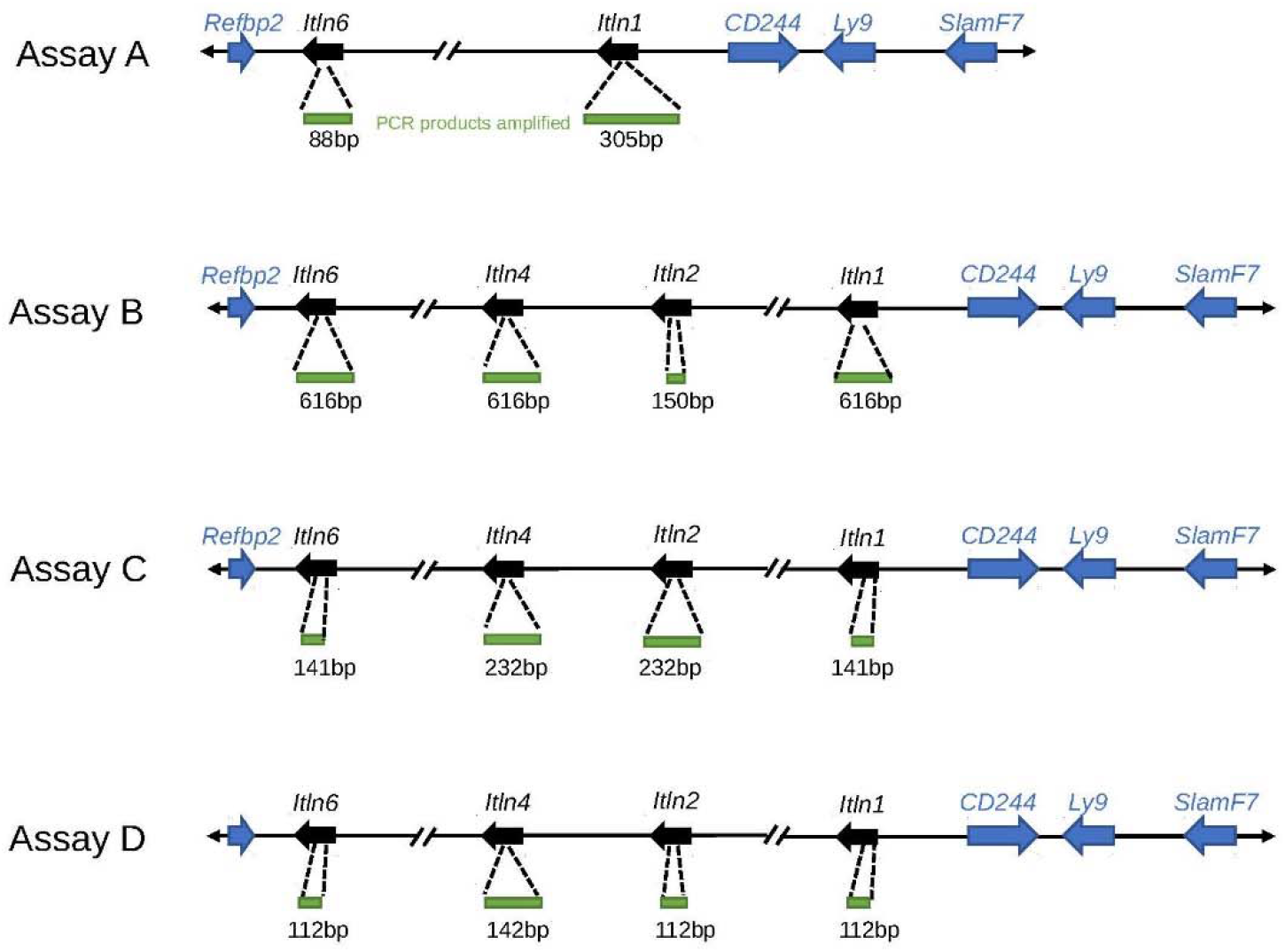
Paralog ratio tests (PRTs) to measure relative copy number of mouse intelectin genes. Four assays (A-D) designed to amplify different sized PCR products (green) using a single primer pair in each assay. Predicted amplicon lengths from the contig of the region previously generated from the 129S7 mouse strain.

**Figure 5.**
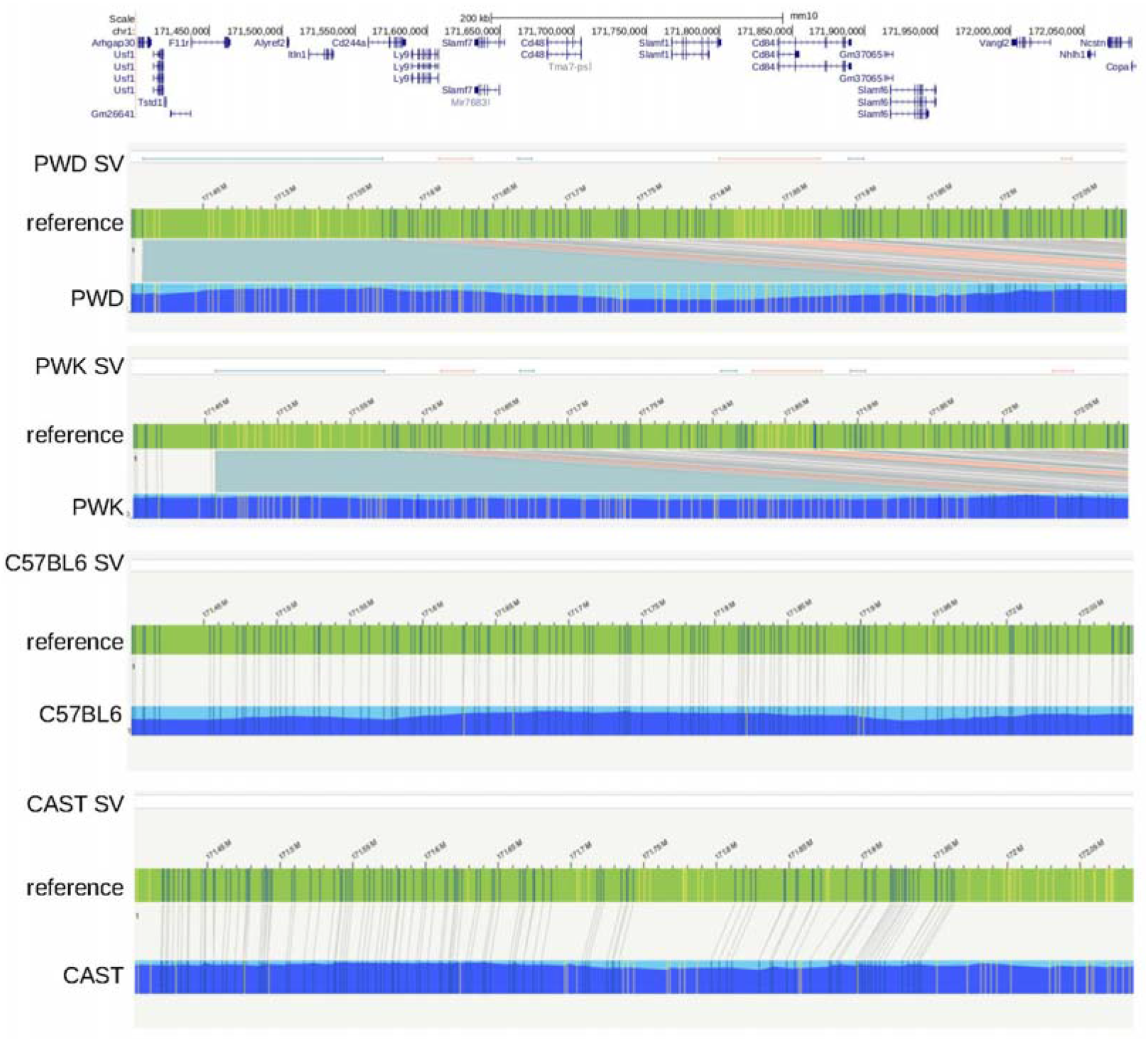
Optical mapping of the intelectin region in laboratory strain mice. Optical mapping results from four strains (PWD, PWK, C57BL/6NCrl, CAST/EiJ) have been aligned against the reference genome (green bars, annotated with genes and long terminal repeats (LTRs). Regions showing structural variation, with respect to the C57BL6/J reference genome, are annotated as SV, green lines showing putative insertions and red lines showing putative deletions. An extra ~400kb of sequence is in both PWK and PWD strains in the *Itln1* region, consistent with the presence of the full complement of intelectin genes. Gene annotations are at the top, and are taken from the UCSC Genome Browser.

It was previously suggested that the genome of the CAST/EiJ strain has a different deletion from that observed in C57BL/6J (8), although neither deletion was characterised. Since CAST/EiJ is a strain derived from wild-caught *Mus musculus castaneus*, we tested the hypothesis that in wild-derived mouse strains other copy number variants could exist, and we made use of previously sequenced mouse strains, including wild-derived mouse strains, to determine CNV across the intelectin region. We analysed 15 laboratory mouse strains that had previously been whole-genome sequenced using Illumina short-read sequencing technology (16,17). Because the alignment and variant analysis of these strains had used the reference C57BL/6NJ genome, and subsequent genome assemblies of a selection of these strains also only assembled *Itln1*, we remapped raw paired-end sequence reads to the previously-assembled 603 kb contig spanning the intelectin region (8). For each sequenced mouse strain, we then plotted normalised sequence read depth in 5kb windows across the assembly. The normalised sequence read depth was not uniform across the region because of variation in the density of repeat-masked sequence, and therefore the number of reads that are mapped, varies across different 5kb windows. However, the noted variation was highly reproducible across different strains known to have the full complement of intelectins. Compared to 129S7/Sv, as expected, sequence from C57BL/6NJ showed a drop in read depth corresponding to the known loss of intelectin genes *Itln2*, *Itln3*, *Itln4*, *Itln5*, and *Itln6*.

The CAST/EiJ strain mouse was found to have a similar size deletion as C57BL/6NJ (Figure 6), which was confirmed using optical mapping (Figure 5). Fine mapping using our paralogue ratio tests and paralogue-specific PCR showed that the CAST/EiJ deletion resulted in loss of *Itln2* and *Itln5*, but not *Itln6*, delineating it from the deletion event found in C57BL/6NJ (Supplementary table 2). The PANCEVO/EiJ strain, derived from *Mus hortulanus*, also showed a deletion involving the same genes as CAST/EiJ. In the SPRET/EiJ mouse strain, derived from *Mus spretus*, a smaller contiguous deletion was found, which deleted *Itln2*, *Itln3* and also *Itln5* but not *Itln1*, *Itln6* or *Itln4* (Figure 6). A similar deletion was found in SKIVE/EiJ (mosaic of *M. m. Musculus* and *M. m. domesticus*) and MOLF/EiJ (*M. m. mollosinus*) strains (Supplementary table 2), suggesting that the deletion occurs across subspecies of *M. musculus*. Several attempts were made to design specific PCR amplifications to precisely determine deletion breakpoints, but these failed due to the highly repetitive nature of the sequence around the breakpoints.

**Figure 6.**
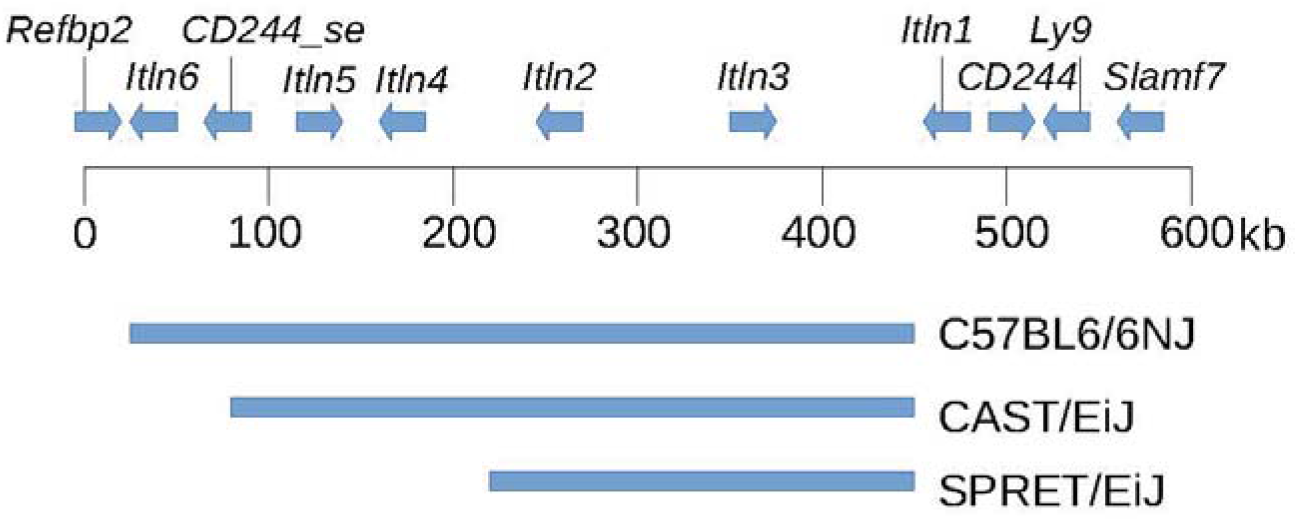
Deletions found by sequence analysis of laboratory strain mice. The region covered by the original 129S7/Sv contig is shown with genes annotated. Extent of deletions found by sequence read depth analysis in laboratory strain mice shown below the size scale.

We extended our analysis from laboratory inbred strains to wild-caught outbred mice. We analysed publicly-available sequence data from 29 *Mus musculus domesticus*, *Mus spretus* and *Mus musculus castaneus* mice (18). A small deletion was discovered in a wild *M. musculus castaneus* mouse from the Himalayas in India (sample H28), and a wild *M. musculus domesticus* mouse from Germany (sample TP51D). This deletion removes both *Itln2* and *Itln3*, and is distinct from the deletions observed in the laboratory strains, suggesting that this deletion is polymorphic across wild mouse subspecies populations. This deletion was confirmed by PRT analysis.

### Mouse intelectin genes *Itln3* and *Itln5* are polymorphic pseudogenes

We analysed our sequencing read alignments from the 15 laboratory mouse strains and 29 wild-caught mice for novel single nucleotide variants in the coding regions of the intelectin genes. Because of the sequence similarity between intelectin genes in this family, care was taken to minimise the risk of incorrectly identifying apparent single nucleotide variants that were in fact differences due to mismapping of sequence reads from similar, paralogous, sequences. Three factors suggest that this was not a problem in our analyses. Firstly, the ability to readily visualise deletions by sequence read depth analysis suggests that extensive mismapping from non-deleted to deleted regions does not occur. Secondly, alignment of sequence from 129S1/SvlmJ strain against the reference derived from the same strain (129S7/Sv) gave no variants across the intelectin region, suggesting no mismapping of reads. Finally, mapping C57BL/6NJ sequence reads to the contig containing all the intelectin genes resulted in only a single, low quality, variant call across the region deleted in C57BL/6J. No variants were annotated in the deleted regions of CAST/EiJ or SPRET/EiJ. Taken together this suggests a very low or non-existent level of sequence mismapping. Nevertheless, all variant calls were visually inspected at the sequence alignment level to check for reads with multiple mismapping sites and for a biologically appropriate ratio of alternative allele counts (i.e. ~0.5 for heterozygotes, and ~1 for homozygotes).

No variants in the coding regions of the intelectin genes were observed in 129S1/SvlmJ, BALB/cJ or C3H/HeJ, suggesting a recent shared origin of this region across these three strains. For the other strains that have a full complement of intelectin genes, between 16 and 31 variants per strain were observed across the intelectin genes. For the strains with a deletion, CAST/EiJ had 12 variants and SPRET/EiJ had 40. We focused on single nucleotide variants that were predicted to introduce or removed stop codons from each gene (Table 1), as these we would expect to have a major effect on biological function. A variant in *Itln5* exon 5 changed a cysteine to a stop codon in DBA/2J, CBA/J, A/J, PWK/PhJ and NZO/HlLtJ, and this premature termination codon is likely to suppress translation of the resulting mRNA by nonsense mediated decay (19). Conversely, in exon 4 of *Itln3*, a stop codon present in the reference sequence is a CAG (Gln) in FVB and WSB strains, suggesting that this gene, although annotated as a pseudogene on the mouse reference genome, has potential for expression of a full-length intelectin in these strains.

**Table 1.** putative functional amino acid variants identified in intelectin genes – (all lab mice strains shown were homozygous for the alternative variant)

| location | Amino acid position | Gene | Ref codon | Alt codon | Population | Allele frequency |
| --- | --- | --- | --- | --- | --- | --- |
| 28777 | Y297F | *Itln6* | TAC<br>Y | TTC<br>F | Germany (Heligoland) | 0.3 |
|  |  |  |  |  | Germany (Cologne-Bonn) | 0.81 |
|  |  |  |  |  | France (Massif Central) | 0.68 |
|  |  |  |  |  | Iran (Ahvaz) | 0.5 |
|  |  |  |  |  | CAST/EiJ, WSB/EiJ, FVB/NJ, NZO/HILtJ, NOD/ShiLtJ, NZB/B1NJ, PWK/PhJ, and A/J | (Inbred strains) |
| 117243 | Q103* | *Itln5* | CAG<br>Q | TAG * | Germany (Cologne-Bonn) | 0.25 |
| 118425 | C235* | *Itln5* | TGT<br>C | TGA * | France (Massif Central) | 0.25 |
|  |  |  |  |  | DBA/2J, CBA/J, A/J, PWK/PhJ, NZO/HILtJ | (Inbred strains) |
| 174162 | T261S | *Itln4* | ACT<br>T | TCT<br>S | NZB/B1NJ | (Inbred strains) |
|  |  |  |  |  | Germany (Cologne-Bonn) | 0.25 |
|  |  |  |  |  | France (Massif Central) | 0.125 |
|  |  |  |  |  | Iran (Ahvaz) | 0.375 |
| 365046 | *143Q | *Itln3* | TAG * | CAG<br>Q | Germany (Cologne-Bonn) | 0.312 |
|  |  |  |  |  | Iran (Ahvaz) | 0.5 |
|  |  |  |  |  | FVB, WSB | (Inbred strains) |
| 463452 | T261S | *Itln1* | ACT<br>T | TCT<br>S | SPRET/EiJ | (Inbred strains) |
|  |  |  |  |  | Germany (Cologne-Bonn) | 0.18 |
|  |  |  |  |  | France (Massif Central) | 0.18 |
|  |  |  |  |  | Iran (Ahvaz) | 0.125 |
| 463445 | H263A | *Itln1* | CAT<br>H | GCT<br>A | Germany (Cologne-Bonn) | 0.18 |
|  |  |  |  |  | France (Massif Central) | 0.18 |
|  |  |  |  |  | Iran (Ahvaz) | 0.125 |
|  |  |  |  |  | FVB, WSB | (Inbred strains) |

As expected for individuals from outbred populations, wild mice show much more single nucleotide variation, with each individual having between 14 to 72 variants. Wild mice populations are polymorphic for both the *Itln5* stop allele and the *Itln3* stop allele seen in certain laboratory strains, suggesting that some laboratory strains inherited these alleles from the wild population. In addition, another *Itln5* early truncation allele, altering a glutamine at position 103 to a stop, is observed in the *M. m. domesticus* population sampled from Cologne in Germany.

As intelectins are calcium-dependent lectins, we also examined variation affecting either the known calcium- or carbohydrate-binding amino acid residues between and across the intelectin family (2,13). An alignment of the mouse and human intelectin genes shows that all seven of the carbohydrate-binding amino acids (residues 243, 244, 260, 262, 263, 274, 288 and 297) are identical between human ITLN1 and mouse Itln1, consistent with glycan-binding biochemical studies for these two proteins (2). However, for other intelectins at least one of these amino acids varies (Figure 7, highlighted in red). For *Itln6*, the carbohydrate-binding tyrosine at position 297 is polymorphic with an alternative non-polar phenylalanine allele (Table 1), potentially altering carbohydrate-binding. Two other amino acid variants in *Itln1* and *Itln4* (T261S and H263A) affect amino acids that may directly (i.e., ligand binding) or indirectly (i.e., configuration of binding pocket) modify glycan binding. In contrast, all ten of the calcium-binding residues are perfectly conserved (Figure 7, highlighted in blue).

**Figure 7.**
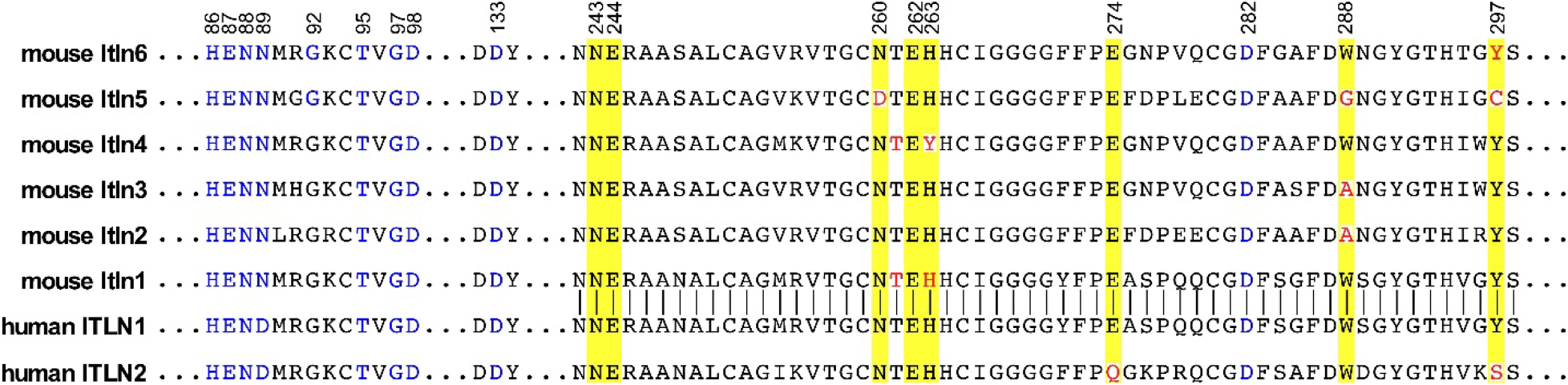
Comparison of the calcium- and glycan-binding regions of mouse and human intelectins. The aligned partial amino acid sequences of human and mice intelectins are shown (amino acid residue numbering from human ITLN1). Amino acids previously shown (2) to mediate calcium coordination are shown in blue and those mediating glycan binding highlighted with yellow. Residues identical to those previously shown to bind carbohydrate in human ITLN1 are shown in black within the yellow background and the polymorphic amino acids are shown in red.

### Mouse Itln1 shows evidence of recent adaptive evolution

Given that mouse intelectin genes clearly have a recent evolutionary history of repeated rounds of duplication followed by deletion and point mutations, we wanted to examine evidence for natural selection at the genes at different times following divergence of the lineages leading to humans and mice. To do this, we used the variation data generated for each gene in wild mice and applied the McDonald-Kreitman test to compare polymorphism within mice, and divergence between mice and other species at synonymous and non-synonymous codon sites (Table 2). In this case, we took each mouse intelectin coding sequence in turn, and compared the polymorphism within that gene to divergence against mouse paralogues, the single rat orthologue, and a human orthologue (*ITLN1*), with the aim of detecting selection at recent timescales. There is evidence of selection of mouse *Itln1* following the burst of duplication that generated multiple intelectins (Table 2). Because of the likelihood of gene conversion events homogenising sequence between recent duplications, a date for the duplication event cannot be reliably estimated from sequence divergence. However, since the rat has only one intelectin gene we assume that these duplication events occurred after rat-mouse divergence between 9-14MYa (20).

**Table 2.**
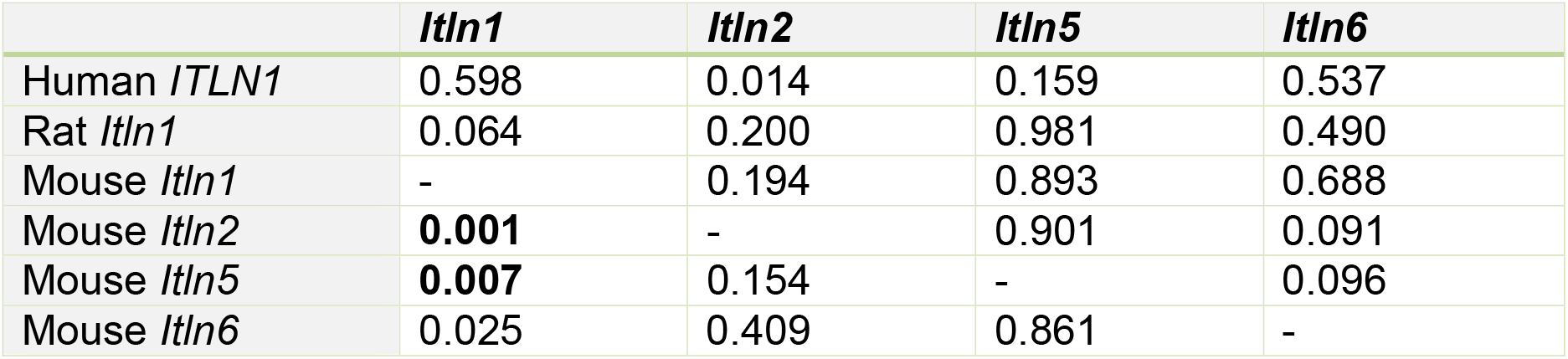
McDonald-Kreitman analysis of mouse *Itln* genes. P values are shown indicating the comparison between variations within the *Itln* gene (columns) against divergence (rows). Statistically significant values with a false discovery rate of <10% are shown in bold.

## Discussion

Intelectins are a family of calcium-dependent lectins that span vertebrate evolution. Human ITLN1 is implicated in several disease states, including inflammatory bowel disease, asthma, and obesity; however, its specific effector functions remain unclear. Mouse models are powerful tools to elucidate the biological function of specific proteins *in vivo*; however, such studies for intelectins are complicated because of the striking, and as yet incompletely characterized strain-specific variation of the intelectin locus resulting in uncertainty regarding which mouse intelectin gene(s) might represent the corresponding orthologue in humans. Herein, we provide evidence that human *ITLN1* has multiple orthologues in mice, whereas human *ITLN2* does not have a corresponding mouse orthologue. In addition, our data reveal key differences within the mouse intelectin gene family, even between common laboratory strains. Together, our findings provide insight into the evolution of mouse intelectins and highlight the need to carefully consider mouse strain when designing and interpreting experiments whose outcome might be dependent upon intelectin biology and/or report intelectin-specific results.

Phylogenetic tree analysis supports that unlike rats and other murids, which have a single intelectin gene, mice have six intelectin genes resulting from a recent burst of duplications. In humans and other primates, an independent duplication event occurring early in (or preceding) primate evolution generated two intelectin genes, *ITLN1* and *ITLN2*. We provide evidence that based on overall protein length and N-terminal sequence of the mature peptide, ITLN2-like proteins are distinguished from ITLN1-like proteins. Our phylogenetic analysis indicates that there is no simple one-to-one relationship between human and mouse intelectin genes. Instead, our data support that all six mouse intelectins are ITLN1-like.

We extensively characterized strain-specific variation of mouse intelectin genes, including analysis of common laboratory strains and wild mice. A previous report demonstrated that the 129S7/Sv laboratory mouse strain has six intelectin genes, whereas the C57BL/6J strain encodes only one due to a 420 kb deletion (5). Using a newly developed collection of paralogue ratio tests we determined that 15 common laboratory mouse strains have the full complement of six intelectin genes. In contrast, all tested sub-strains of the C57 line have a single family member, *Itln1*. Using PCR primers designed to amplify across the deletion breakpoint in C57BL/6J, sequence data from the PCR products confirmed that the breakpoint was identical in these sub-strains, including C57 parent lines: C57L/J, C57BR/cdJ, and C58/J. Optical mapping further supported these assertions. These results imply that the deletion event occurred prior to human/researcher-driven mouse inbreeding. Similar analysis indicated that a deletion in CAST/EiJ and PANCEVO/EiJ mice removes *Itln2* and *Itln5,* and a deletion in SPRET/EiJ, SKIVE/EiJ, and MOLF/EiJ strains removes *Itln2*, *Itln3* and *Itln5,* indicating that these deletions are not identical to the C57BL/6 deletion.

We detected no novel single nucleotide variants in the coding regions of the intelectin genes the genome sequences of 129S1/SvlmJ, BALB/cJ or C3H/HeJ mice, suggesting a recent shared origin of this region across these three strains. However, for the other laboratory strains, including those that have the full complement of six intelectin genes, we observed between 16 and 31 coding-region variants per strain, similar to inbred CAST/EiJ (12 variants) and SPRET/EiJ (40 variants), which possess locus deletions. As expected, we observed a higher frequency of single nucleotide variants in wild-caught mice (between 14 to 72 variants per individual). Of the single nucleotide variants predicted to have profound consequences, a premature stop codon (C235*) in exon 5 of *Itln5* was observed in five lab strains (DBA/2J, CBA/J, A/J, PWK/PhJ, and NZO/HlLtJ) and a wild-caught mouse, and a glutamine codon replaced the premature stop codon in exon 4 of *Itln3* (*143Q) of the reference genome, in two lab strains (FVB and WSB) and two wild-caught mice.

Complementary evidence supports the concept that Itln1 is the functional homologue of human ITLN1 (5). We highlight here that both genes encode a preproprotein of identical length (313 aa), and that the residues responsible for calcium coordination (10 residues) and those mediating carbohydrate binding (8 residues) are identical in the two orthologous proteins. Prior biochemical studies confirm that glycan-recognition selectivity of Itln1 was analogous to its human counterpart, however subtle differences in binding selectivity, kinetics, and affinity were reported (2,5). Our data also show that mouse *Itln1* is expressed in the small intestine confirming initial reports (4).

Our findings suggest that adaptive evolution has occurred in mouse *Itln1* during the last 9-14 million years, since repeated rounds of duplication resulted in expansion of the intelectin locus. The amino acid changes in Itln1 adjacent to the carbohydrate-binding residues may have resulted in subtle alteration of affinity or spectrum of carbohydrate-binding during mouse evolution. Moreover, tissue specificity outlined here and/or potential inducibility may define different functions of individual mouse intelectins (21). Other differences between species are that human ITLN1 exists as a disulfide-linked trimer, whereas mouse Itln1 lacks cysteines required for intermolecular disulfide-linked trimer formation (5), and that within the intestine, mouse *Itln1* is expressed in Paneth cells, whereas human ITLN1 is expressed in goblet cells. Nevertheless, a strong argument can be made for mouse Itln1 to have essentially similar functions in the gut as human ITLN1.

The relationships and functions of the other mouse intelectins are less clear. As noted, our data support that all six mouse intelectins are *ITLN1*-like, with no mouse orthologue of human *ITLN2*. Mouse *Itln2*, despite its numerical designation, is not the orthologue of human *ITLN2*. Of the six encoded intelectins, only *Itln1* and *Itln6* were found to be constitutively expressed at high levels, where *Itln1* is present in the small intestine and *Itln6* is present in the colon. It is interesting that like *Itln1*, *Itln6* shares amino acid identity at the eight positions mediating glycan binding, although a conservative polymorphism at Y297F was noted in some strains. We speculate that perhaps differences in gene regulation and/or protein secretion in development and/or under environmental stress will explain the distinguishing anatomic patterns of expression.

From the data presented here, other mouse intelectin genes have variant residues at one or more key amino acids that govern carbohydrate binding, have polymorphic early truncating codons and/or have polymorphic large deletions. Combined with the lack of evidence for adaptive evolution at these other intelectin genes, this suggests a lack of critical function, and that these genes are, or are in the process of becoming, pseudogenes. Therefore, we may be witnessing a snapshot of an evolutionary process, likely common in other immunity gene families, called birth-and-death evolution (22). Future work requires focus on identifying any novel functions of these mouse intelectin genes, in particular those that contribute to a high expression level in the uterus, which does hint at a possible function. If any gene does encode a fully functional protein, it is likely that polymorphic variation within wild mouse populations and between laboratory strains will have functional consequences.

This study also emphasises the importance of examining natural variation for gene regions that are absent in the reference genome of the organism under study (23). While now being addressed in humans using long-read sequencing and efforts by consortia like the telomere-to-telomere consortium, analysis in model or domestic animals lags behind. Moreover, our study highlights the careful considerations required by researchers investigating innate immunity genes in mice, where strain and sub-strain variations can complicate interpretations of gene function (24–26).

## Methods

### Mouse husbandry and tissue analysis

The Institutional Animal Care and Use Committee at the University of California, Davis, approved all procedures involving live animals and methods of euthanasia; experiments were performed following AVMA guidelines and with strict adherence to IACUC-approved protocols. Briefly, animals were deeply anesthetised with a cocktail of ketamine and xylazine (100/mg/kg and 10 mg/kg, respectively) prior to euthanasia. Tissue samples were dissected immediately after mice were euthanized and submerged in RNAlater™(Ambion Inc, Austin, TX). The RNAlater specimen tubes were incubated with gentle rocking overnight at 4°C, and then stored long-term at −20 °C.

### DNA samples and extraction

Genomic DNA was isolated and purified using the QIAamp DNA minikit (Qiagen, Germantown, MD) according to the manufacturer’s protocol. Isolated DNA was quantified by ultraviolet absorbance spectroscopy (260 nm) using a NanoDrop spectrophotometer (Thermo Scientific/NanoDrop Products, Wilmington, DE).

### Phylogenetic analysis

Intelectin coding sequences for the phylogenetic tree were retrieved from the list of orthologues and paralogues of human *ITLN1* curated by Ensembl (release 98). Mouse *Itln* sequences were identified from the full length 129 contig accession number HM370554.

Evolutionary analyses were conducted in MEGA X (27). For the protein tree, amino acid sequences were inferred from coding DNA sequences and aligned using ClustalW, and the phylogenetic tree inferred by maximum likelihood and JTT matrix-based model, Atlantic salmon (*Salmo salar*) *Itln1* was used as an outgroup. The tree with the highest log likelihoo is shown. Initial tree(s) for the heuristic search were obtained automatically by applying Neighbor-Join and BioNJ algorithms to a matrix of pairwise distances estimated using a JTT model, and then selecting the topology with superior log likelihood value. A discrete Gamma distribution was used to model evolutionary rate differences among sites (5 categories (+G, parameter = 0.4683)). The rate variation model allowed for some sites to be evolutionarily invariable, but no sites were found to fit in that category of the model. The tree is drawn to scale, with branch lengths measured in the number of substitutions per site. Following bootstrap analysis, the percentage of trees in which the associated taxa clustered together is shown next to the branches.

### RNA isolation and expression analysis

The general procedures for RNA isolation and synthesis of cDNA were previously described by our group (28,29). Briefly, the RNAlater solution was decanted and the tissue was homogenized in guanidine thiocyanate buffer. Total RNA was isolated using cesium chloride gradient ultracentrifugation, quantified using ultraviolet absorption spectrometry at 260 nm, and reverse transcribed to cDNA using an oligo-(dT)12-18 primer. The single-stranded cDNA product was purified using the Qiagen PCR purification kit (Qiagen, Valencia, CA), and diluted to 10 ng/µl based on the input concentration of total RNA. Real-time PCR was performed using as templates the cDNA from experimental tissues and gene-specific plasmids as external standards for quantification. The primer pairs for qPCR were: *Itln1* forward 5’- ACCGCACCTTCACTGGCTTC-3’, *Itln1* reverse 5’- CCAACACTTTCCTTCTCCGTATTTC-3’, and *Actb* forward 5’-GGCTGTATTCCCCTCCATCG-3’, reverse 5’- CCAGTTGGTAACAATGCCATGT-3’. Absolute quantification of specific mRNA from tissue was determined by extrapolation of the detection threshold (crossing point) to the crossing point for gene-specific external plasmid standard analyzed within each run. Reproducibility assessments of this approach were previously reported (28). For analysis across multiple strains, data were analyzed using the delta CT method normalizing gene expression to *Actb* in each sample. A negative control reaction that omitted template cDNA was included with each set of reactions to check for possible cross-contamination.

For sequence analysis of RT-PCR products, intestinal cDNA (distal small intestine and colon) from 129S2/SvPasCrl mice was used as a template in a PCR reaction using oligonucleotide primers whose sequences correspond to regions of identical sequence in mRNA of all six paralogues of intelectin (m129ItlnCom-4s 5’- GCCTCAGCAGAGAAAGGTTCC-3’ and m129ItlnCom-287a 5’GAAGGTCTGGTAGATGACACCATTC-3’). Primers were designed using MacVector Software (MacVector, Apex, NC) and synthesized by Invitrogen Life Technologies (Invitrogen, Carlsbad, CA). The PCR reactions were initiated by denaturation of the DNA template at 95°C for 10 min followed by 45 cycles consisting of 95°C for 15 sec, a −1°C per two-cycle ‘touchdown’ annealing temperature for 5 sec (i.e., 65°C to 58°C), and 10 sec at 72°C. The PCR product was purified by passage through a PCR cleanup column following the manufacturer’s protocol (Qiagen). The purified sample was then sequenced using an Illumina platform (Genewiz, South Plainfield, NJ). Sequence data was filtered to remove reads that failed base-calling quality checks, and the reads with identical sequence were assigned to a particular intelectin gene by comparison to a reference sequence.

### DNA Sequence analysis

All mouse genome coordinates are using the GRCm38 assembly. The coding sequence for human *ITLN1*, human *ITLN2* and Rat *Itln1* were retrieved from the NCBI-nucleotide database (https://www.ncbi.nlm.nih.gov/nuccore) with accession numbers AB036706.1, AY065973.1 and XM_017598901 respectively. The coding sequence for each mouse *Itln* gene was retrieved from the BAC contig of the 129S7 intelectin locus from GenBank with accession number (HM370554) (https://www.ncbi.nlm.nih.gov/nuccore/HM370554).

Whole genome sequencing (WGS) data for 29 wild-caught mice were accessed at the European Nucleotide Archive (ENA; http://www.ebi.ac.uk/ena/) under accession number (PRJEB9450).These 29 wild-caught mice were sequenced by Illumina Hiseq 2000 at the Max Planck institute (Harr *et al.*, 2016). The 15 laboratory mouse strains were sequenced by the Wellcome Trust Sanger institute and accessed at the European Nucleotide Archive (ENA; http://www.ebi.ac.uk/ena/) (16) (Supplementary table 3).

The quality of the raw fastq files was assessed using FastQC v 0.11.5, followed by adapter removal using Cutadapt v 01.11 (30). Following mapping of the processed fastq files to the repeat-masked BAC contig (accession number HM370554) using BWA-MEM v 0.7.15 (31), the resulting SAM file was converted to a BAM file and sorted and indexed using SAMTools v 1.3.2 (32). Multiple bam files from the same mouse sample were merged using SAMtools, and labelled as a single new unified read group by Picard v2.1 (33). The resulting bam file was subjected to local realignment using GATK 3.6 and 3.8 and duplicate removal using Picard v 2.1, resulting in the final single bam file for each mouse.

### Paralogue Ratio Test

The paralogue ratio test (PRT) is a form of quantitative PCR that uses shared PCR primers to minimise differences in amplification kinetics between cases and controls (34,35). We designed four paralogue ratio tests (PRTs, assays A-D) to examine relative copy numbers of intelectin genes in mice by designing a primer pair specific to a subset of intelectin genes, ensuring at least 5 mismatches between the primer and non-amplified intelectins. PCR primers were 5’-TATTCCTGTCTCAGCTCCTAG-3’, 5’- GTCACAGGTAAAKCCAGAAGG-3’ for assay A, 5’-TGTAAAYCCCTCCTTGACTCC-3’, 5’-GATAAATGRCCCAGTMCTTGCC-3’ for assay B, 5’-CACACGTACACCTTCCTG-3’, 5’-GAGTAGTCTGCYKTATTTCAG-3’ for assay C and 5’- GTAATGYTGACTTYGGCCTTC-3’, 5’-GTTGAGCWTGGCAGATTGT-3’ for assay D with IUPAC codes K Y, M and R indicating that the primers were synthesized with a mix of the two indicated nucleotides at that position.

PCR amplification was in a final volume of 10 ul, which included 5-10 ng genomic DNA, 0.5U Taq DNA polymerase and a final concentration of 1mM primer in 1x KAPA Buffer A (a Tris-ammonium sulphate buffer with a final concentration of 1.5mM MgCl_2_).

Thermal cycling was one cycle of initial denaturation at 94°C for 2 minutes, followed by 35 cycles of denaturation at 94°C for 30 seconds, then PRT1 and PRT2 were annealed at 60°C and PRT3 and PRT4 were annealed at 62°C. All assays were annealed for 30 seconds; this was then followed by an extension step for 30 seconds at 72°C. Lastly, one extra extension step was carried out at 72°C for 5 minutes. PCR amplification products from different intelectin genes were detected based on amplicon size using standard 2% agarose gel electrophoresis, ethidium bromide staining and visualization under UV light.

### Calling variation from sequence alignment files

Across the whole region (603274bp) of the *Itln* locus, the number of reads mapping to non-overlapping 5kb windows was calculated using SAMTools (v1.3.2) (32). In each single window, reads were counted, normalized to average read count and plotted to visualize gain or loss. Single nucleotide variation for the exonic regions for each individual mouse from the bam file was called using FreeBayes (v1.1) (36), and converted to vcf format using VCFtools (v0.1.14) (33) on a minimum quality score of 30 and a minimum depth of 15. All vcf files are available at https://doi.org/10.25392/leicester.data.13679035.v1.

### Optical mapping

We generated optical maps across the whole-genome of four different mice, from two mouse subspecies. C57BL/6J (B6) and C57BL6Crl (B6N) of *Mus musculus domesticus,* CAST/EiJ of *M. musculus castaneus* and PWD/Ph (PWD) and PWK/Ph (PWK) of *M. m. musculus* origin. First megabase-scale high molecular weight DNA was extracted according to the Saphyr Bionano Prep Animal Tissue DNA Isolation Soft Tissue Protocol (Document Number: 30077; Document Revision: B). Briefly, cell nuclei were isolated from splenic tissue and embedded in agarose plugs. DNA in plugs was purified with Proteinase K and RNAse, then high molecular weight (HMW) genomic DNA was extracted from the agarose plugs using agarase, and purified by drop dialysis. HMW DNA was resuspended overnight before quantification with the Qubit BR dsDNA assay, then kept at 4°C until labelling.

We performed the Bionano Direct Labelling and Staining (DLS) protocol (Document Number: 30024 Revision:I) on 750ng of DNA from each sample using direct labelling enzyme (DLE-1) to label all its recognition sites (CTTAAG). After an initial clean-up step, the labelled HMW DNA was pre-stained, homogenized, and quantified with the Qubit HS dsDNA assay, before using an appropriate amount of backbone stain YOYO-1. The molecules were then imaged using the Bionano Saphyr System (Bionano Genomics, San Diego). The resulting de-novo optical maps were then generated and mapped against an in-silico optical map of the mouse genome reference sequence.

## Acknowledgements

The Genotype-Tissue Expression (GTEx) Project was supported by the <u>Common Fund</u> of the Office of the Director of the National Institutes of Health, and by NCI, NHGRI, NHLBI, NIDA, NIMH, and NINDS. We acknowledge research grant support from the National Institutes of Health (U01AI125926 [Mucosal Immunology Study Team] and R37AI32738).

## Author contributions

Conceptualisation – FA, EBN, EJH and CLB

Investigation – all authors

Resources – EJH, CLB, LO-H and BL

Supervision – EJH, CLB, LO-H and BL

Visualisation – FA, EBN, LO-H and EJH

Writing – original draft – EJH, CB, EBN and FA

Writing – review and editing – all authors

---

**Supplementary figure 1.**
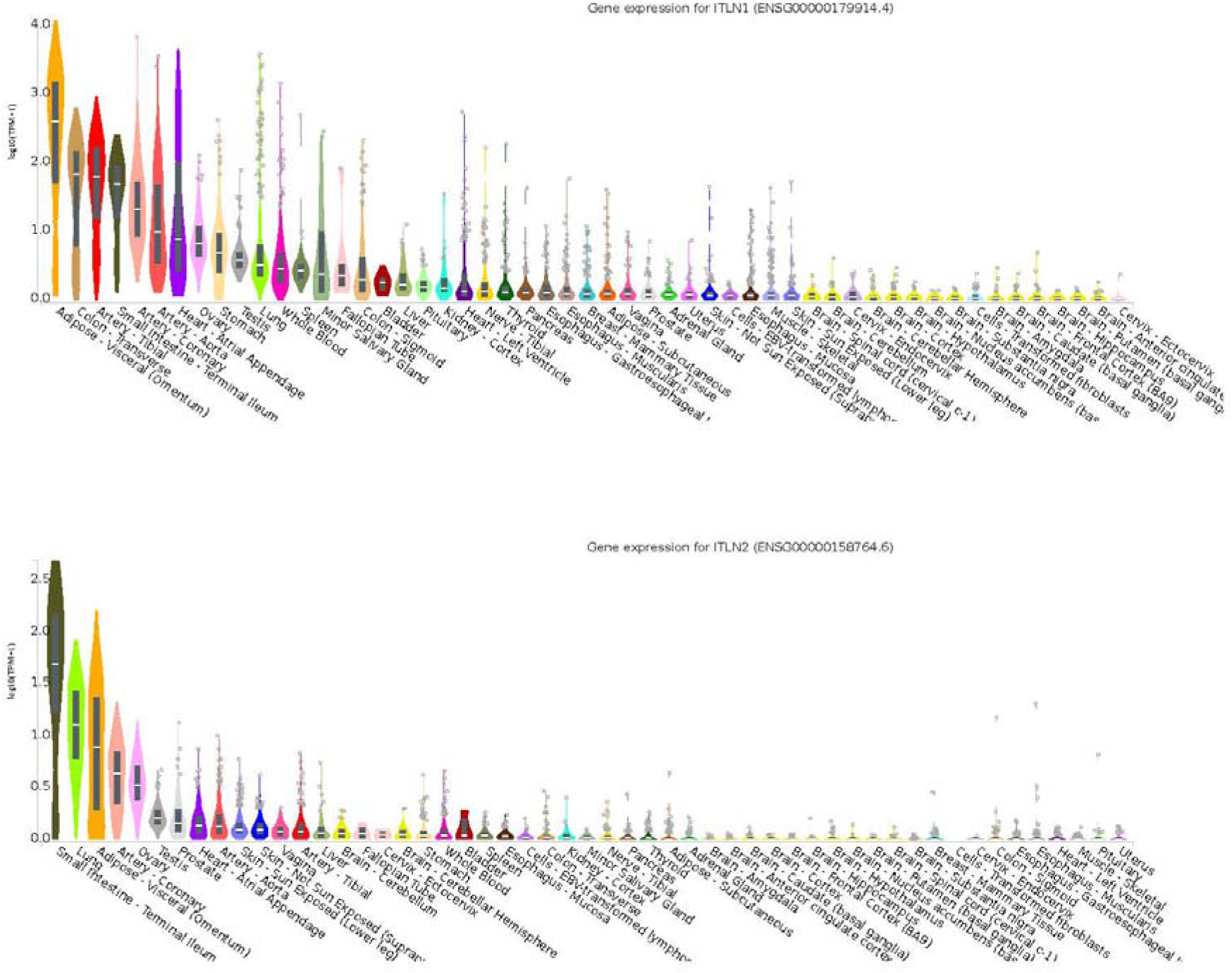
*ITLN1* and *ITLN2* gene expression patterns in humans Expression levels measured by RNAseq over a number of different human individuals for 53 tissues, ordered by median expression level. Data from Gtex database, plotted using GTEX browser.

**Supplementary figure 2.**
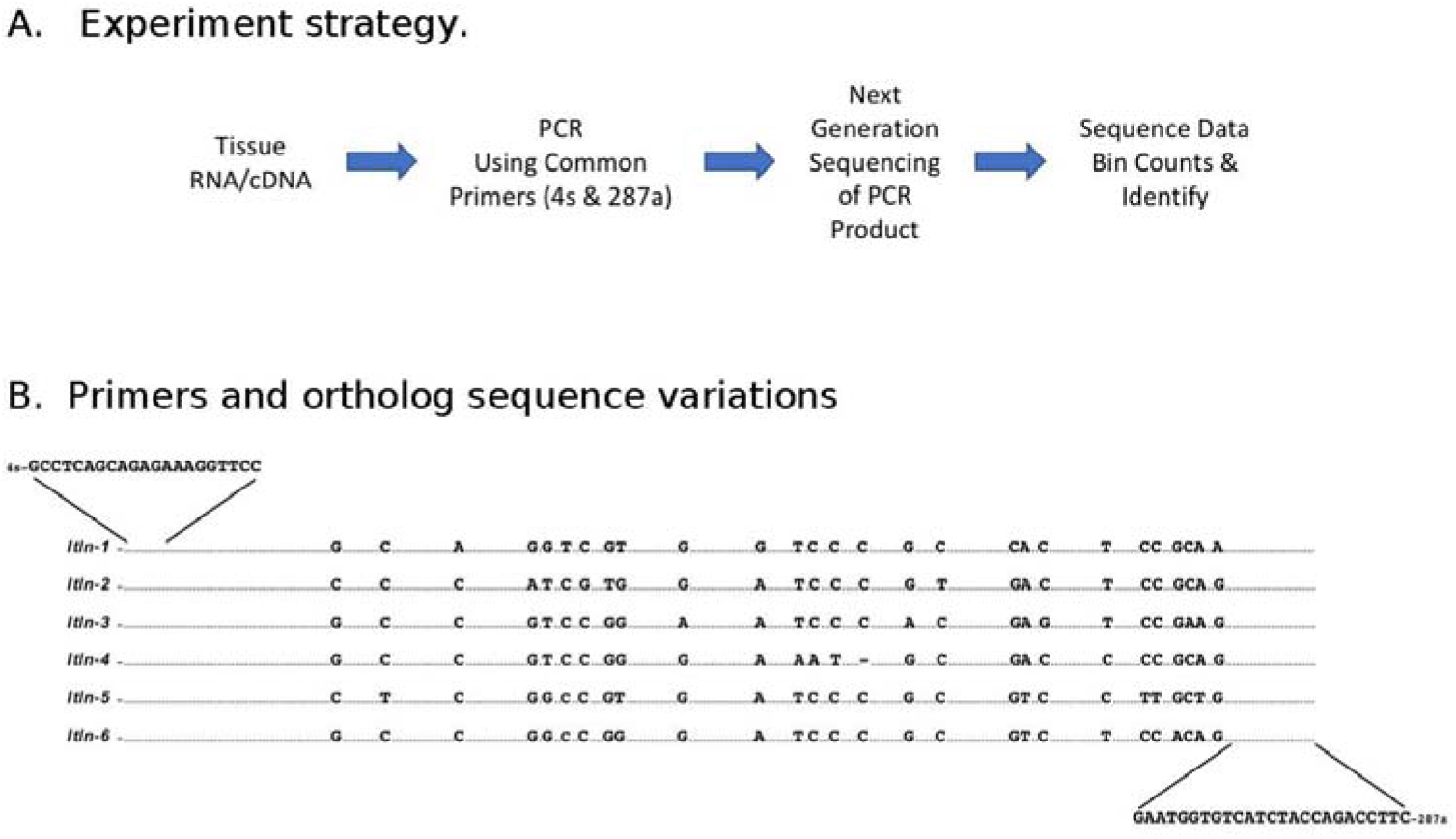
High throughput sequencing strategy to measure relative expression abundance of individual mouse intelectins. A. Outline of strategy. B. Sequence alignment of PCR product (283 nt) showing nucleotides that differ among the six mouse intelectin genes.

**Supplementary Table 1.** Relative mRNA abundance (%) of mouse intelectin paralogues in small intestine and colon of 129S2/SvPasCrl, as determine by high-throughput sequencing.

| Gene | Sequence Bin Count |  |
| --- | --- | --- |
|  | Small Intestine | Colon |
| ***Itln1*** | 112,357<br>(99.83%) | 0 |
| ***Itln2*** | 246<br>(0.22%) | 48<br>(0.06%) |
| ***Itln3*** | 0 | 0 |
| ***Itln4*** | 0 | 0 |
| ***Itln5*** | 0 | 0 |
| ***Itln6*** | 168<br>(0.15%) | 82,566<br>(99.94%) |

**Supplementary Table 2.** Presence of particular intelectin genes in mouse strains determined by PRT. P=present, A=absent, ND= not determined.

| Subgenus, Species, & Subspecies | Strain/Substrain | Vendor:ID | *Itln1* | *Itln2* | *Itln4* | *Itln6* |
| --- | --- | --- | --- | --- | --- | --- |
| *Mus musculus* – mosaic:<br>*domesticus/musculus* | C57BL/6NCrl | CRL:574 | P | A | A | A |
|  | C57BL/6NTac | TB: B6 | P | A | A | A |
|  | C57BL/6NJ | JL: 005304 | P | A | A | A |
|  | C57BL/6J | JL: 000664 | P | A | A | A |
|  | C57BL/10J | JL: 000665 | P | A | A | A |
|  | C57BL/10SnJ | JL: 000666 | P | A | A | A |
|  | C57BLKS/J | JL: 000662 | P | A | A | A |
|  | C57L/J | JL: 000668 | P | A | A | A |
|  | C57BR/cdJ | JL: 000667 | P | A | A | A |
|  | C58/J | JL: 000669 | P | A | A | A |
|  | B6N-Tyrc-Brd/BrdCrCrI | CRL: 493 | P | A | A | A |
|  | C57BL/6NTac-<br>*tm1Arte*<br>*Tyr* | TB: 11971 | P | A | A | A |
|  | B6(Cg)-*Tyr*<sup>*c-2J*</sup>/J | JL: | P | A | A | A |
|  |  | 000058 |  |  |  |  |
|  | 129S1/SvImJ | JL:<br>002448 | P | P | P | P |
|  | 129S2/SvPasCrl | CRL: 476 | P | P | P | P |
|  | 129S4/SvJaeJ | JL:<br>009104 | P | P | P | P |
|  | 129S6/SvEvTac | TB:<br>129SVE | P | P | P | P |
|  | 129S8/SvEvNimrJ | JL:<br>012809 | P | P | P | P |
|  | 129X1/SvJ | JL:<br>000691 | P | P | P | P |
|  | 129P3/J | JL:<br>000689 | P | P | P | P |
|  | BALB/cAnNCrl | CRL: 547 | P | P | P | P |
|  | BALB/cAnNTac | TB: BALB | P | P | P | P |
|  | BALB/cJ | JL:<br>000651 | P | P | P | P |
|  | BALB/cByJ | JL:<br>001026 | P | P | P | P |
|  | DBA/1J | JL:<br>000670 | P | P | P | P |
|  | DBA/2J | JL:<br>000671 | P | P | P | P |
|  | CBA/J | JL:<br>000659 | P | P | P | P |
|  | C3H/HeJ | JL:<br>000659 | P | P | P | P |
|  | A/J | JL:<br>000646 | P | P | P | P |
|  | AKR/J | JL:<br>000648 | P | P | P | P |
|  | FVB/NJ | JL:<br>001800 | P | P | P | P |
|  | NOD/shiLtJ | JL:<br>001976 | P | P | P | P |
|  | SJL/J | JL:<br>000686 | P | P | P | P |
|  | SWR/J | JL:<br>000689 | P | P | P | P |
|  | CD-1 IGS | CRL: 022 | P | P | P | P |
|  | CrI:CFW(SW) | CRL: 024 | P | P | P | P |
|  | CrI:NIHBL(S) | CRL: 492 | P | P | P | P |
| ***Mus musculus domesticus*** | LEWES/EiJ | JL:<br>002798 | ND | ND | ND | ND |
|  | PERC/EiJ | JL:<br>001307 | ND | ND | ND | ND |
|  | WSB/EiJ | JL:<br>001145 | P | P | P | P |
| ***Mus musculus musculus*** | SKIVE/EiJ | JL:<br>001393 | P | A | P | P |
| ***Mus musculus musculus*** | MOLF/EiJ | JL:<br>000550 | P | A | P | P |
|  | PWD/PhJ | JL:<br>004660 | P | P | P | P |
|  | PWK/PhJ | JL:<br>003715 | P | P | P | P |
|  | SF/CamEiJ | JL:<br>000280 | P | P | P | P |
| ***Mus caroli*** | CAROLI/EiJ | JL:<br>000926 | P | P | P | P |
| ***Mus musculus castaneus*** | CAST/EiJ | JL:<br>000735 | P | A | A | P |
| ***Mus hortulanus*** | PANCEVO/EiJ | JL:<br>001384 | P | A | P | P |
| ***Mus pahari*** | *Mus pahari*/EiJ | JL:<br>002655 | ND | ND | ND | ND |
| ***Mus spretus*** | SPRET/EiJ | JL:<br>001146 | P | A | P | P |

**Supplementary table 3.** Mouse sequences used. Accession numbers for raw laboratory mouse strains WGS data that were sequenced by the Wellcome Trust Sanger institute. Accessed at the European Nucleotide Archive

| **Sample ID** | **Accession Number** | **Species** |
| --- | --- | --- |
| 129S1/SvImJ | ERS076385 | *Mus musculus* |
| A/J | ERS075416, ERS138733,<br>ERS212195 | *Mus musculus* |
| BALB/cJ | ERS076386 | *Mus musculus* |
| C3H/HeJ | ERS076383 | *Mus musculus* |
| C57BL/6NJ | ERS076384 | *Mus musculus* |
| CAST/EiJ | ERS076381 | *Mus musculus*<br>*castaneus* |
| CBA/J | ERS076379 | *Mus musculus* |
| DBA/2J | ERS075663 | *Mus musculus* |
| FVB/NJ | ERP000687 | *Mus musculus* |
| NOD/ShiLtJ | ERS076389 | *Mus musculus* |
| NZB/B1NJ | ERS661331 | *Mus musculus* |
| NZO/HILtJ | ERS076387 | *Mus musculus* |
| PWK/PhJ | ERS076378 | *Mus musculus musculus* |
| SPRET/EiJ | ERS076388 | *Mus spretus* |
| WSB/EiJ | ERS076380 | *Mus musculus*<br>*domesticus* |

